# *Anopheles* spp. distribution and climatological niche modeling to predict malaria potential along bioclimatic envelope gradients in South Coast of West Java landscape

**DOI:** 10.1101/2021.04.18.440313

**Authors:** Andri Wibowo

## Abstract

Malaria remains a major public health problem mainly in particular South East Asian countries. As malaria transmission and *Anopheles* spp. continues to spread, control interventions should emphasize on the ability to define potential areas that can favor *Anopheles* spp. distribution. Then there is an urgent need to use novel approach capable to predict potential spatial patterns of *Anopheles* spp. and delineate malaria potential hotspots for better environmental health planning and management. Here, this study modeled *Anopheles* spp. potential distribution as a function of 15 bioclimatic variables using Species Distribution Modeling (SDM) in South Coast of West Java Province spans over 20 km from West to East. Findings of this study show that bioclimatic variables and SDM can be used to predict *Anopheles* spp. habitat suitability, suggesting the possibility of developing models for malaria early warning based on habitat suitability model. The resulting model shows that the potential distributions of *Anopheles* spp. encompassed areas from West to Central parts of the coasts, with Central parts were the most potential prevalence areas of *Anopheles* spp. considering this area has higher precipitation. The less potential prevalence areas of *Anopheles* spp. were observed in the East parts of the coast. The model also shows that inland areas adjacent to the settlements were more potential in comparison to the areas near coast and in the beach. Land cover conditions dominated by cropland, herbaceous wetland, and inundated land were also influencing the *Anopheles* spp. potential distribution.

## INTRODUCTION

Malaria is still considered as a potential zoonotic disease in many developing countries where it is endemic through the economic and health burden it imposes on those countries. Malaria also remains one of the most serious public health problems associated with high morbidity and mortality in the world. This disesase is a climate-sensitive protozoan disease driven by parasites of *Plasmodium* genus and transmitted among humans through bites of infected female *Anopheles* mosquitoes. It was estimated that 3.2 billion people remain at risk of malaria world-wide resulting in 438000 deaths annually. In spite of significant progress made in reducing malaria morbidity and mortality through intensification of malaria and *Anopheles* control programs, most developing countries are still at risk of malaria. For instance, WHO estimated 214 million new cases of malaria were reported in 2015.

Indonesia is one of tropical countries in South East Asia that is still threatened by malaria. According to WHO data in 2010 there were 544470 cases of malaria in Indonesia, where in 2009 there were 1.1 million clinical cases and in years 2010 it increased again to 1.8 million cases. In 2011, the number of malaria cases was 256592 people and 1.3 million cases of suspected malaria with blood samples were examined showed an Annual Parasite Insidence (API) of 1.75 per thousand population. This means that for every 1000 inhabitants in endemic areas there are 2 people infected by malaria. Malaria transmission in Indonesia continues to occur and national health research reports show that until 2011 there were 374 districts that were endemic to malaria.

According to the Guidelines (WHO 2007) on the elimination of residual foci of malaria transmission, malaria foci are determined by the spatial presence of parasite, host, and vector populations. Spatial based entomological surveillance is an important tool to determine the receptivity to malaria in malarious areas, defined as areas in which transmission of malaria is occurring or has occurred during the preceding three years. In addition, investigation and modeling of the vulnerability to infection based on the host *Anophele*s presence is an important part of surveillance efforts in malarious areas to determine the magnitude of the malariogenic potential. Therefore, there is urgent need to use geospatial based distribution modeling tools such as geographic information based system to detect spatial patterns and forecast the potential distribution of *Anopheles* and malaria, and delineate disease hot spots for better planning and management.

In Indonesia, one of vulnerable areas is located in the South Coast of West Java Province (Hakim et al. 2018, Nababan & Umniyati 2018.). This province is knowing having the highest population in comparison to other 33 provinces. The South coast is also dominated by wetland and microclimate that may be suitable and favorable for the *Anopheles* spp. to breed and increase the risks of malaria transmission to the community nearby.

## MATERIALS AND METHODS

### Study area

This study was conducted in several locations across 60 km coast line of South coast of West Java. Those locations geographically were in longitude of 106.0-108.0 East and latitude of 6.0-8.0 South (Figure 1). The lowland landscapes and land covers of South coast of West Java were dominated by the combinations of cropland, herbaceous wetland, and inundated land.

**Figure 1.**
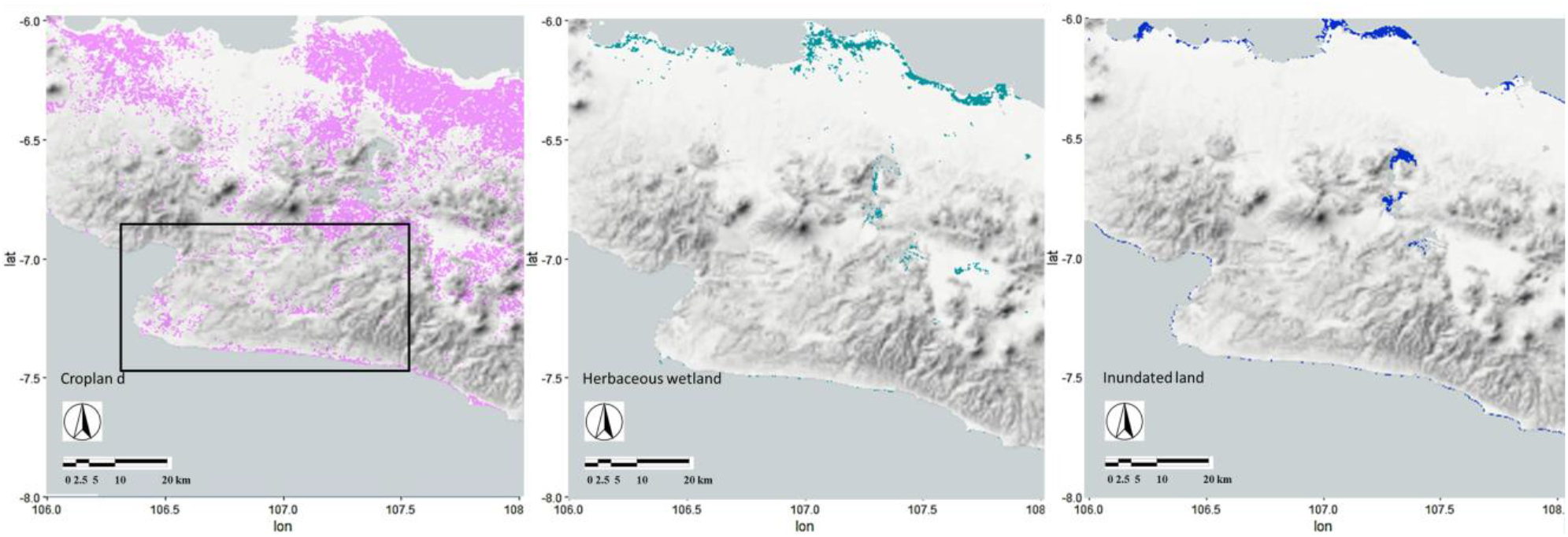
Studied locations (rectangle) (lon: 106.0-108.0 E, lat: 6.0-8.0 S), land covers (from left to right: cropland, herbaceous wetland, and inundated land), and landscapes in South Coast of West Java Province, Indonesia.

### Bioclimatic envelope

The species distribution modeling performed in this study was based on the species preference on climate at local scale and bioclimatic envelope concept following Rose (1998). Bioclimatic envelope modeling is used to describe the present and future distribution of ecological elements, whether individual species or entire life zones, based on suitable climate conditions. The model’s development and subsequent application is supported by niche theory coined by Vandermeer (1972), Austin (2002), and Leibold (1995) that describes the climatic niche as a functional or conceptual space defined on multiple axes of climatic variables. The climatic niche is one aspect of an organism’s or ecosystem’s fundamental niche, and it is assumed to remain static and does not take dispersal ability or evolutionary adaptation into consideration when extrapolating from current distributions to future potential. Pearson and Dawson (2003) stated that bioclimatic envelopes are appropriately employed at regional scales where climate has a dominant influence on species and geographically calibrated bioclimatic envelopes are an inherently conservative tool for habitat modeling. It allows firm identification of some known acceptable bioclimatic variables, even if the definition of all acceptable climates is incomplete.

According to Kadmon et al. (2003), this novel modeling strategy is ideally suited to presence-only data, a characteristic of most conservation targets. In this study, there are 15 bioclimatic variables variables that have been modeling using WorldClim’s database with high spatial resolution global weather and climate data. Those bioclimatic variables including annual mean temperature, mean diurnal range, isothermality, temperature seasonality, maximum temperature in warmest month, minimum temperature in coldest month, temperature annual range, mean temperature in wettest quarter, mean temperature in driest quarter, mean temperature in warmest quarter, mean temperature in coldest quarter, annual precipitation, precipitation in wettest month, precipitation in driest month, precipitation seasonality, precipitation in wettest quarter, precipitation in driest quarter, precipitation in warmest quarter, and precipitation in coldest quarter.

In developing bioclimatic envelope using WorldClim (Fick & Hijmans 2017, Marchi et al. 2019), weather station data were obtained from multiple sources. Station data were checked for correspondence between their reported elevation and the elevation obtained from a global elevation raster data. Stations with large deviations (>several 100 m) between reported and actual elevation were mapped and evaluated relative to available geographic information and neighbouring station data. Average temperature was calculated as the mean of maximum and minimum of tabulated station-wise monthly temperatures.

### Species distribution modeling (SDM)

Species distribution modeling for *Anopheles* spp. was following Akpan et al. (2018) and Hanafi-Bojd et al. (2018). First, the *Anopheles* spp. presence was recorded based on real time mosquito surveys (Sugiarto et al. 2009) in designated study area. Observed *Anopheles* spp. was recorded for its geocoordinate positions including longitude and latitude. The geographically presences of *Anopheles* spp. then were mapped into thematic layers of study areas. Species distribution models or ecological niche models (SDMs), also known as bioclimatic envelope models, a most widely used approach for predicting suitable habitats for different species, was developed from the bioclimatic data including monthly temperature and precipitation values in order to generate more biologically meaningful variables. Those bioclimatic variables represent annual trends, seasonality, and extreme or limiting environmental factors. In this study, the bioclimatic data were downloaded from the WorldClim database. This dataset is available as an individual raster spanning the inhabited continents and presented as latitude/longitude coordinates in WGS84. In this study, all 15 bioclimatic variables with a spatial resolution of 1 km^2^ were retrieved in designated South Coast of West Java.

All selected *Anopheles* spp. presences and bioclimatic variables for modeling were converted to ASCII format for modeling in the next step by SDM that take a list of species presence locations as input, often called presence-only data, as well as a set of bioclimatic variables functioning as predictors including precipitation and temperature variables. The final output is a thematic layer with classifications of *Anopheles* spp. environmental suitability ranges from high to low.

## RESULTS AND DISCUSSION

### *Anopheles* spp. spatial occurrences

Figure 2 presents the realtime *Anopheles* spp. occurrences and distribution along the coast. In total there were 22 locations were confirmed for *Anopheles* spp. occurrences. The occurrences were in agreement with the presences of cropland, herbaceous wetland, and inundated land that were favorable microhabitats for *Anopheles* spp. This finding and area preferences were in corroborated with the presences of *Anopheles* spp. in other locations mainly in coastal environments. Leaua (2013) and Suwito et al. (2010) confirmed that wetlands adjacent to settlement and croplands were commonly invested with *Anopheles* spp. The cropland and forest fragment in the study areas have the potential to retain rainfall water (Albuquerque et al. 2018). This ground water is gradually released into the rivers favoring the existence of permanent water bodies in the region as the inundated lands were common in here. The studied areas were mostly located in low elevations and the extensive areas of lowlands can accumulate water from rivers and water ways in higher elevation due to the low slope of the terrain. As a result, areas in coast with low elevation contain water bodies that are potential larval habitats for *Anopheles* spp.

**Figure 2.**
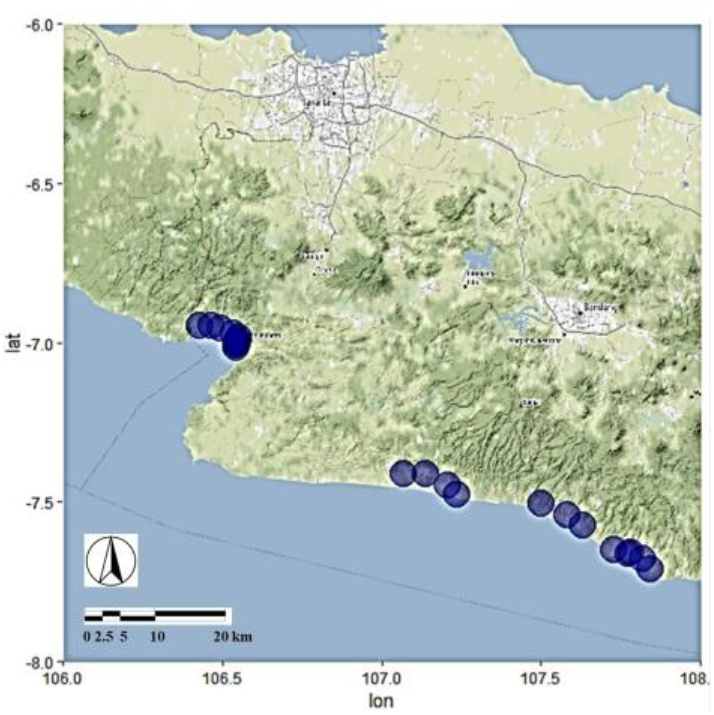
*Anopheles* spp. occurrences (blue dots) in South Coast of West Java.

### Bioclimatic variables

Figure 3 presents the 15 bioclimatic variables retrieved from WorldClim and modeled in studied areas. The bioclimates of South Coast of West Java were characterized by high temperature within the ranges of 26-32 °C and high rainfall in some areas. This bioclimatic characteristrics are typical climate of tropical coast. The occurrences of *Anopheles* spp. as a function of bioclimatic variables in particular annual mean temperature and precipitation can be seen in Figure 4. It can be seen that in the coast, the *Anopheles* spp. occurences were following coastal climates of 26 °C. The occurrence variations can be seen in bioclimatic variables of precipitations. In the West parts of the coast, *Anopheles* spp. was occurred in coastal areas with higher precipitations (400 mm/year) in comparison to other areas. While in East parts, coastal areas with lower precipitations equals to 250 mm/year were observed favorable for *Anopheles* spp. In this coast, the precipitations were decreasing from mountainous areas in the West to the lowland in the East. In tropical ecosystems, *Anopheles* spp. occurences were known having positive correlation with precipitations. This species can occur in areas with precipitation ranges of 15.6-211.5 mm/year with the highest precipitation of 439 mm (Tulak et al. 2018). Whereas Kazwaini & Willa (2015) observed that the presences of aqautic habitats are more correlated with the *Anopheles* spp. occurences.

**Figure 3.**
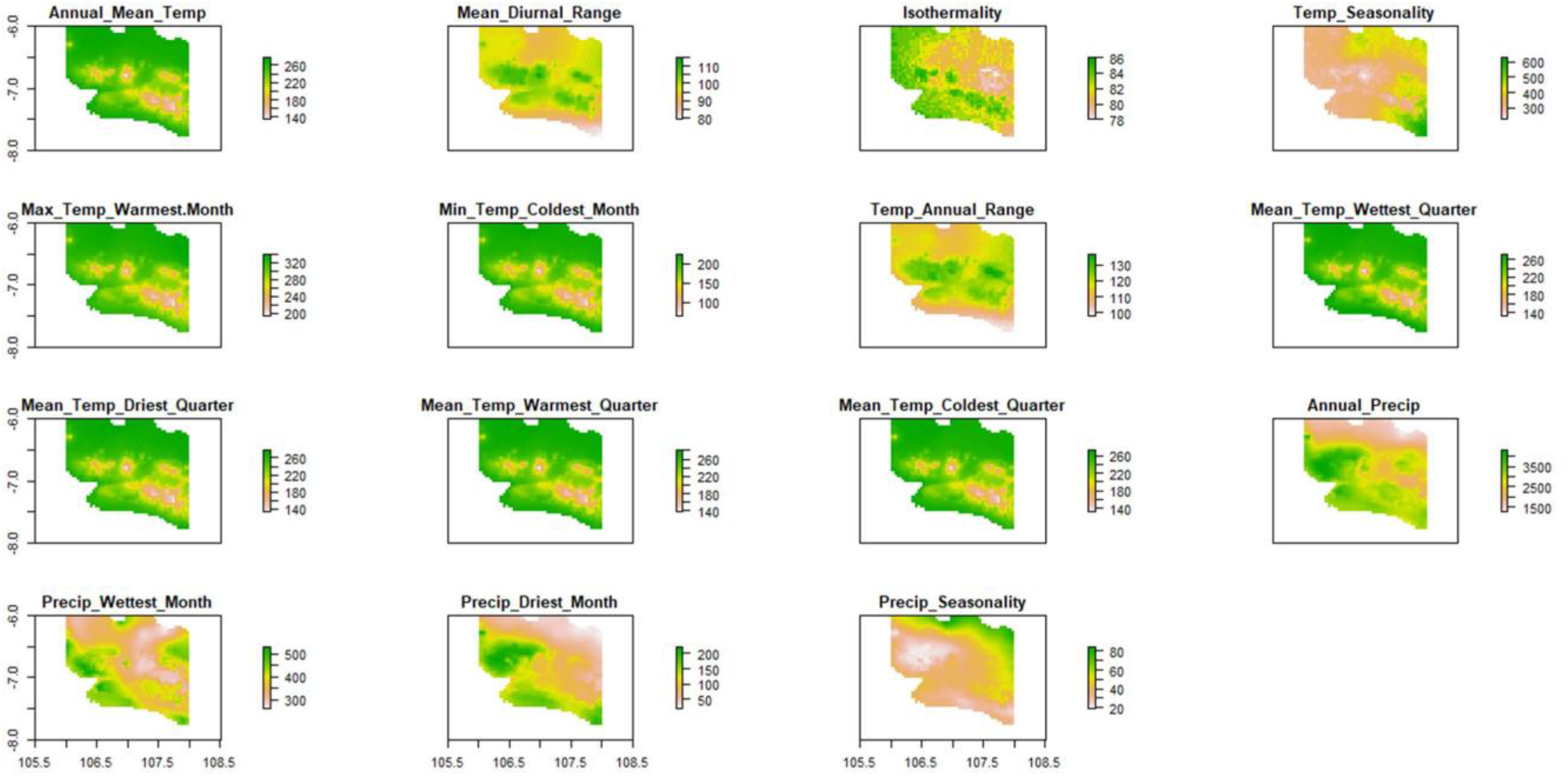
Bioclimatic variables retrieved from WorldClim and modeled in South Coast of West Java.

**Figure 4.**
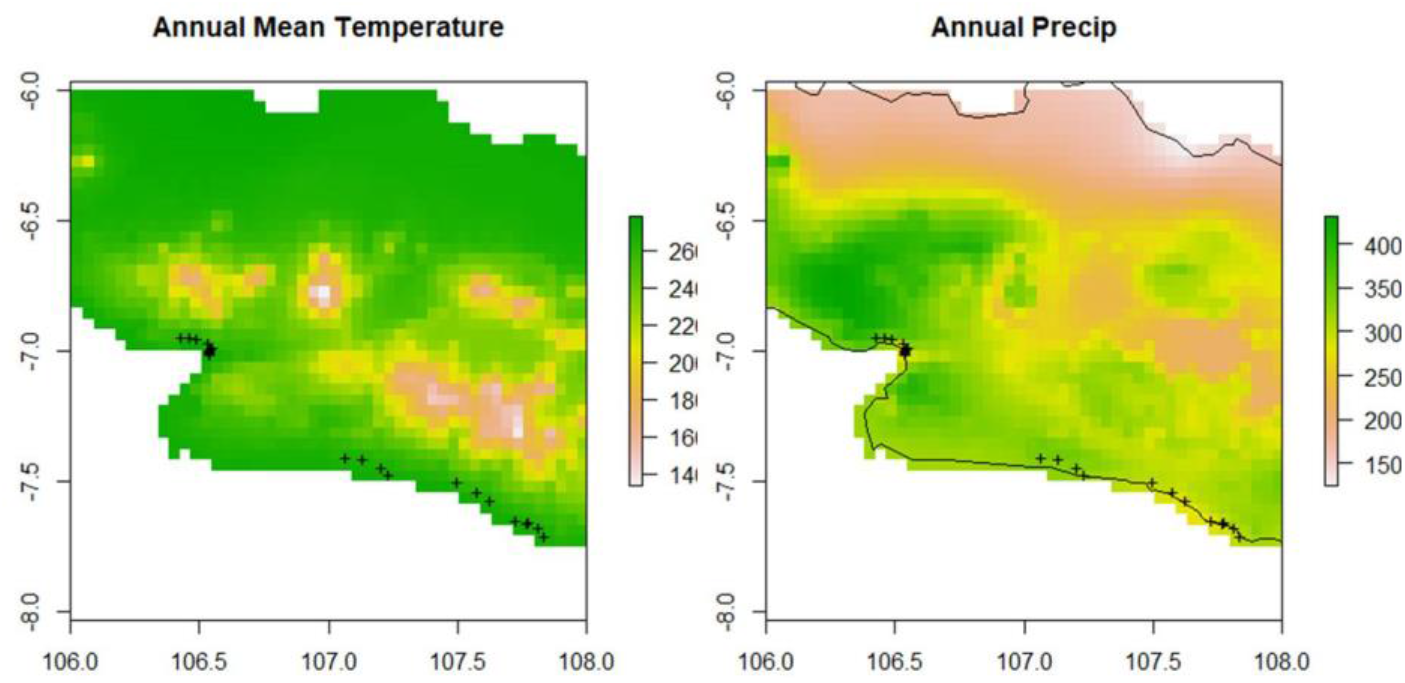
*Anopheles* spp. occurrences (+ marks) along annual mean temperature (°C) and annual precipitation (mm/year) in South Coast of West Java.

This study analyzed the preference and occurrence of *Anopheles* spp. based on bioclimatic variables retreived from WorldClim database and this is with an agreement with other studies. Remote sensing based bioclimatic variable models offer an opportunity to improve species disatributioon modeling and prediction world-wide. Waltari et al. (2014) stated that the weather station-based WorldClim data set has been the primary source of temperature and precipitation information used in correlative species distribution models. In this study, *Anopheles* spp. was presence in coastal areas characterized with low elevation, high temperature, and precipitation. In agreement with the findings in this study, Albuquerque et al. (2018) confirmed that the areas classified as very high in term of *Anopheles* spp. presnece are characterized by the highest temperatures and precipitation levels and are mostly located in the lowland regions (0–20 m).

### Species distribution modeling of *Anopheles* spp

Figure 5 showed the model of *Anopheles* spp. potential distribution in the South Coast of West Java and this also indicates coastal areas with potential malaria considering *Anopheles* spp. is the host for this zoonotic disease. The resulting model shows that the potential distributions of *Anopheles* spp. encompassed areas from West to Central parts of the coasts, with Central parts were the most potential prevalence areas of *Anopheles* spp. These potential areas are spanning over 20 km from longitude of 106.4 E to 107.3 E. While the less potential prevalence areas of *Anopheles* spp. were observed in the East parts of the coast. Small potential prevalence areas in East parts were related to the lower annual precipitation in the East. For comparison,West and Central parts were having higher precipitations and those were followed by more larger areas that were more suitable for *Anopheles* spp. In West and Central parts, the *Anopheles* spp. was predicted distributed in areas beyond *Anopheles* spp. actual occurrences as can be seen from longitude of 106.3 E to 106.7 E. Even it is predicted that *Anopheles* spp. can occur in the areas where *Anopheles* spp. was absent according to actual surveys. Possible occurrence of a species beyond its actual presence record is possible considering the adjacent areas were having suitable bioclimatic variables and ranges that can favor the species to inhabit those areas. The model also shows that inland areas adjacent to the settlements were more potential in comparison to the areas near coast and in the beach. This finding then warns the settlements near the coast to be more aware on the potential *Anopheles* spp. occurrence and possible malaria incidences.

**Figure 5.**
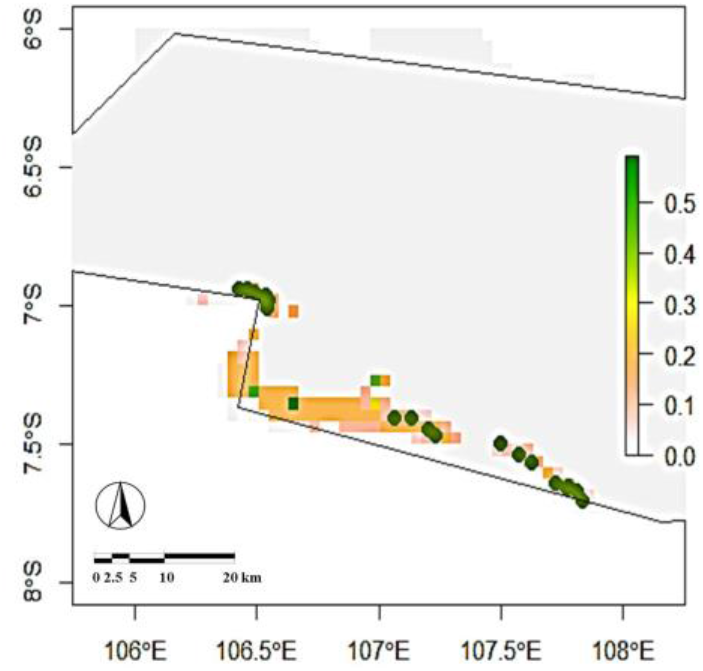
*Anopheles* spp. actual occurrences (green dots) and predicted *Anopheles* spp. potential distribution scales (ranging from low 0.0 to medium 0.25 and high 0.5) according SDM in South Coast of West Java.

The only available SDM study on *Anopheles* spp. and malaria was available in study by Gwitira et al. (2018) and it is limited in the Africa regions. While information on *Anopheles* spp. potential distribution in Asia regions mainly in Indonesia is still lacking. Then this study is the first that provides empirical evidences of *Anopheles* spp. SDM in South East Asian regions. SDM is known as versatile model and approach that have been used to predict potential distribution of species. It is very useful to anticipate the spread of species that can has negative impacts including *Lycorma delicatula* (Hemiptera: Fulgoridae) an invasive insect (Namgung et al. 2020), Tephritid pests invasions (Godefroid et al. 2015), and talaromycosis, an invasive mycosis (Wei et al. 2021).

